# Re-purposing Ac/Ds transgenic system for CRISPR/dCas9 modulation of enhancers and non-coding RNAs in zebrafish

**DOI:** 10.1101/450684

**Authors:** Vanessa Chong-Morrison, Filipa C. Simões, Upeka Senanayake, Dervla S. Carroll, Paul R. Riley, Tatjana Sauka-Spengler

**Affiliations:** University of Oxford, Weatherall Institute of Molecular Medicine, Radcliffe Department of Medicine, Oxford OX3 9DS, United Kingdom; University of Oxford, Department of Physiology, Anatomy and Genetics, Oxford OX1 3PT, United Kingdom; University College London, Department of Cell & Developmental Biology, London WC1E 6BT, United Kingdom

**Keywords:** zebrafish, maize, Ac/Ds, CRISPR, enhancer, non-coding

## Abstract

Due to its genetic amenability coupled with recent advances in genome editing, the zebrafish serves as an excellent model to examine the function of both coding and non-coding elements. Recently, the non-coding genome has gained prominence due to its critical role in development and disease. Here, we have re-purposed the Ac/Ds maize transposition system to reliably screen and efficiently characterise zebrafish enhancers, with or without germline propagation. We further utilised the system to stably express guide RNAs in microinjected embryos enabling tissue-specific CRISPR/dCas9-interference (CRISPR*i*) knockdown of lncRNA and enhancer activity without disrupting the underlying genetic sequence. Our study highlights the utility of Ac/Ds transposition for transient epigenome modulation of non-coding elements in zebrafish.

**Summary statement:** We adapted the Ac/Ds transposition system, which enables continuous expression of guide RNAs for CRISPR/dCas9 perturbation, to examine the function of non-coding RNAs and enhancer elements in zebrafish.

## Introduction

The non-coding genome has risen to prominence in recent years following successive studies highlighting its numerous roles in development and disease (Krijger and de Laat, 2016, Engreitz et al., 2016). The genome is populated by *cis*-regulatory elements called enhancers, which are active in a tissue-specific fashion (Rickels and Shilatifard, 2018) and probing their functional relevance requires inactivation in specific cell types and at distinct times. This is particularly important for key developmental regulators often deployed in well-defined spatiotemporal sequences, and thus likely to employ specific enhancers for such activity. Long non-coding RNAs, or lncRNAs, are defined as transcripts > 200bp with no known protein product (Quinn and Chang, 2015). Unlike in the case of their protein-coding counterparts, functional studies to dissect lncRNA function can be challenging due to limiting factors such as their under-characterisation, low expression levels, short transcript length and rapid degradation. Furthermore, antisense lncRNAs that overlap protein-coding loci but are transcribed on the opposite strand are often found within important developmental loci (Bassett et al., 2014). Uncoupling their function from that of their cognate genes represents a major obstacle in the experimental design of relevant knockout studies.

*In vitro* methods for interrogation of non-coding RNAs and enhancers are useful but may not recapitulate results obtained from *in vivo* transgenic/knockout models, which, on the other hand, are often time-consuming. In this study, we have sought to bridge this gap by developing an efficient and flexible molecular toolkit to functionally assay non-coding elements in the zebrafish using transient and quantifiable *in vivo* approaches. The toolkit was based on re-purposing the maize transposon system first identified by Barbara McClintock in the late 1940s (McClintock, 1950). Molecular characterisation of the system led to identification of the components required for transposition to occur - two Ds (Dissociation) genetic elements and an Ac (Activator) transposase (Fedoroff et al., 1983). Buoyed by this important finding, Ac/Ds elements were employed to facilitate the integration of a reporter construct into the zebrafish genome with high efficiency, leading to remarkable germline transmission rates (Emelyanov et al., 2006). This approach was the integration method of choice used to generate transgenic zebrafish lines for the chemical-inducible LexPR transactivation system (Emelyanov and Parinov, 2008, Kenyon et al., 2018), as well as for a systematic mutagenesis gene-trapping screen (Quach et al., 2015). While these studies demonstrated the utility of Ac/Ds as an efficient method for propagation of transgenes through the germline, they overlooked its strong potential for transient expression of DNA elements in F_0_ embryos, which is a current limitation of the zebrafish model. Several other genome integration methods currently exist in vertebrates (Kawakami, 2007, Vrljicak et al., 2016). In particular in the zebrafish, Tol2-mediated genomic transposition is an established approach for somatic and germline integration of DNA constructs and is almost always the method-of-choice for generating transgenic reporter lines. For transient DNA integration experiments, however, Tol2-mediated transposition often produces variable results with a high rate of F_0_ embryos displaying non-specific background and/or mosaic expression. The analysis of exogenous gene expression and testing of enhancer/reporter activity therefore often relies on the generation of F_1_ offspring carrying relevant constructs. In this study, we expanded the use of Ac/Ds-mediated integration in zebrafish to test and validate putative *cis*-regulatory elements in transient, as an alternative to the Tol2-integration-based approach. This redirects the focus of previous zebrafish Ac/Ds-integration studies from germline propagation efficiency to characterisation of its utility for somatic integration-based experiments.

To efficiently probe non-coding element function in F_0_ embryos in zebrafish, we have used tissue-specific epigenome engineering. Other knockdown approaches in injected F_0_ zebrafish embryos currently exist, such as morpholino-mediated obstruction of protein synthesis or RNA-splicing, or somatic gene editing using TALENs or CRISPR/Cas9. However, these approaches lack spatiotemporal specificity and, as a result, render it difficult to distinguish between a cell-specific phenotype and secondary effects resulting from ubiquitous knockdown of the gene-of-interest. CRISPR/dCas9-based interference (CRISPR*i*) utilises nuclease-deficient Cas9 (dCas9) fused to transcriptional regulator domains which are targeted to specific genomic regions using “guide RNAs” (sgRNAs). To inhibit transcription, dCas9 can be targeted to transcription start sites (TSS) of genes to inhibit RNA Polymerase II by steric hindrance (Qi et al., 2013), or fused to effector domains such as Kruppel-associated box (KRAB) (Gilbert et al., 2013) or four concatenated mSin3 repressive domains (SID4x) (Konermann et al., 2013) to induce chromatin changes inhibitive of transcription. Crucially, as CRISPR*i* complexes are directed to the TSS of target genes, this allows the perturbation of gene expression without modifying the endogenous locus sequence, making it a well-suited tool to study the function of non-coding genes (Thakore et al., 2015, Konermann et al., 2015, Liu et al., 2016, Williams et al., 2018). Similarly, ectopic activation of genes can be achieved by CRISPR*a* in specific cell types using the VP64 activator domain (Mali et al., 2013). The caveat to using these approaches in the zebrafish embryo is the need for extended expression of sgRNAs, which is limited by the current gold-standard of injecting *in vitro*-transcribed sgRNAs. In the absence of Cas9 protein, uncapped and non-polyadenylated sgRNAs degrade quickly (Mir et al., 2018). Therefore such experiments can only be performed at the 1-cell stage and either have to be analysed during the first 24-48 hours of development or sgRNA expression needs to be propagated to the germline. Here, in addition to building zebrafish BAC transgenic lines expressing CRISPR*i*/*a* effectors dCas9-SID4x and dCas9-VP64 in a tissue-specific fashion, we also generated and employed Ac/Ds-integrated DNA constructs to deliver small guide RNAs constitutively in order to transiently modulate the activity of non-coding elements in transgenic F_0_ embryos *in vivo*.

Our Ac/Ds toolkit broadens the potential of the zebrafish embryo for rapid studies of non-coding genomic elements. By harnessing its reliability and efficiency for somatic integration, the toolkit can be robustly used either in transient and/or to complement transgenic-based methods.

## Results and Discussion

### Ac/Ds transposition is more effective than Tol2 for transient integration of transgenes in F_0_ zebrafish embryos

As a first step in re-purposing the maize Ac/Ds transposition system for zebrafish, we generated a new enhancer/reporter construct (pVC-Ds-E1b:eGFP-Ds) for efficient and reproducible *in vivo* testing of enhancer activity. The construct consists of the eGFP expression cassette under the control of the zebrafish E1b minimal promoter (Becker et al., 2016) flanked by Ds elements for integration into the genome (Emelyanov et al., 2006, Emelyanov and Parinov, 2008), and features a multiple cloning site for cloning enhancer sequences to test upstream of the minimal promoter (Fig.1A). We compared the efficiency and efficacy of Ac/Ds- and Tol2-mediated integration (Kawakami, 2004) by transiently expressing a previously predicted zebrafish enhancer for *pax3a* with strong activity (Trinh et al., 2017, Gavriouchkina et al., 2017) (“*pax3a* enhancer”). Ac/Ds and Tol2 versions of the *pax3a* enhancer were microinjected into Gt(FoxD3:mCherry)^ct110aR^ transgenic embryos (Hochgreb-Häagele and Bronner, 2013) together with 24 pg Ac or 50-80 pg Tol2 mRNA, respectively. We observed a clean and tissue-specific pattern of eGFP expression in neural crest cells at the neural plate border with 30 pg of the Ac/Ds vector, a result that could only be recapitulated with 150 pg of Tol2 vector (Fig.1A), indicating higher somatic integration efficiency by Ac/Ds as previously reported (Vrljicak et al., 2016). To quantify this observation, we collected embryos injected with 30 pg of the Ac/Ds or Tol2*pax3a* enhancer for GFP immunohistochemistry (IHC) together with Hoechst-staining of nuclei, followed by confocal imaging and Imaris image analysis (Supp.Fig.1, Materials & Methods). Ac/Ds-injected embryos demonstrated up to 9-fold increase in GFP^+^/Hoechst^+^ signal compared to Tol2-injected embryos (p=0.001) (Fig.1B). The lower load requirement for Ac/Ds (30 pg per embryo) to yield efficient integration also resulted in much lower mosaicity within a clutch of injected embryos (64 to 95% of injected embryos with distinguishable expression pattern, compared to < 40% for Tol2) and therefore provided an efficient binary yes/no tool for transient screening of enhancer activity. To assess the limitations of Ac/Ds-integration in F_0_, we next compared Ac/Ds and Tol2 versions of a previously predicted enhancer for *sox10* with weak activity (scattered labelling of few neural crest cells with either method) (Trinh et al., 2017, Gavriouchkina et al., 2017) (“*sox10* enhancer”) using the same quantification approach (Supp.Fig.2). Tol2-injected embryos demonstrated more mosaic intra-clutch GFP expression pattern compared to Ac/Ds-injected embryos, with a higher proportion of embryos without any specific detectable GFP^+^ cells that were excluded from quantification. Conversely, although weak “*sox10* enhancer” showed smaller number of GFP^+^ cells per embryo, Ac/Ds-integration approach allowed us to detect reproducible and consistent neural crest-specific activity of this enhancer. While Ac/Ds- and Tol2-mediated integration of the ‘weak’ enhancer resulted in comparable GFP^+^/Hoechst^+^ signal, a much higher variability between embryos was observed by the Tol2-mediated approach (Fig.1C).

**Figure 1.**
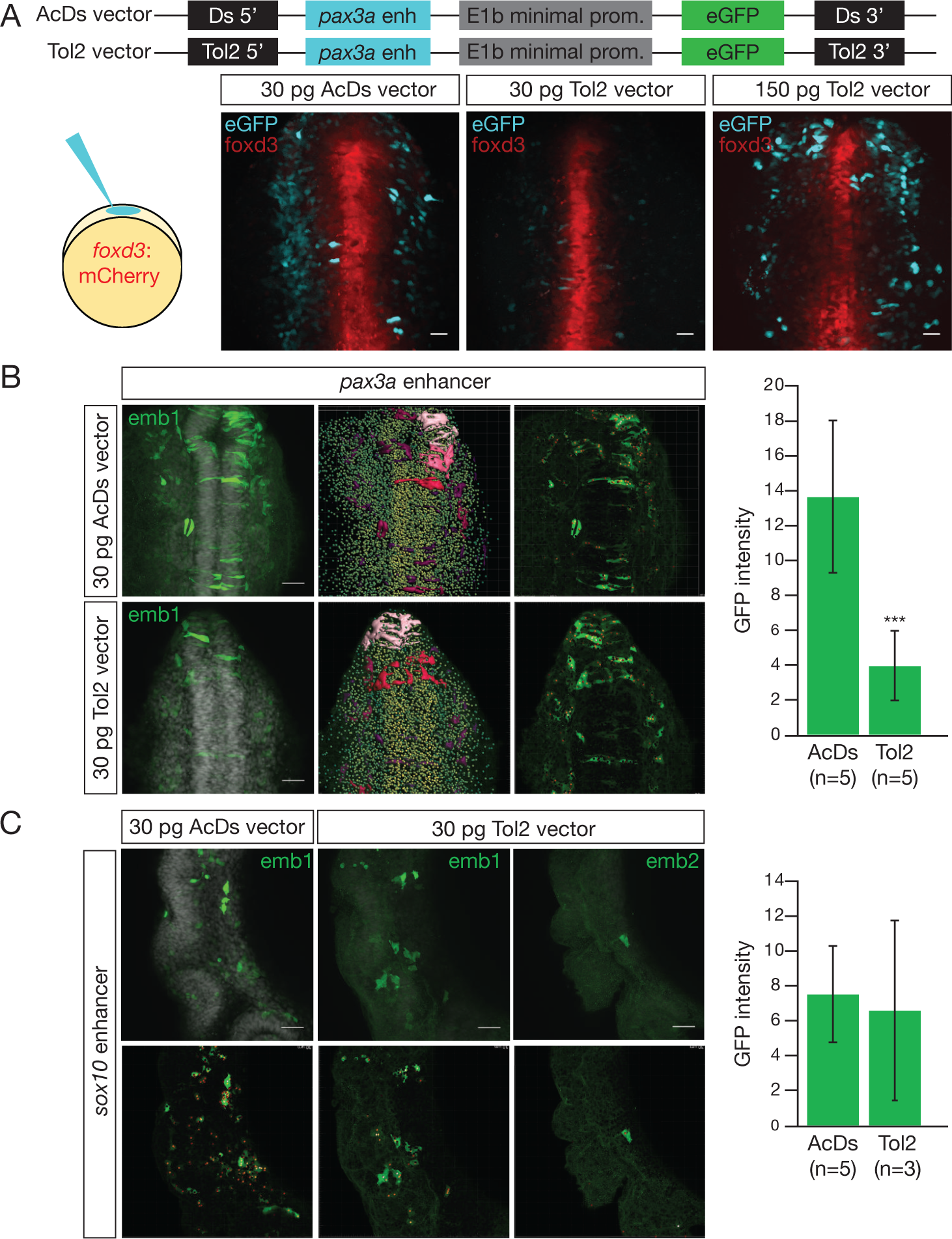
Ac/Ds integration is more effective than Tol2 in F_0_ embryos. **A** Schematic of constructs containing a neural crest *pax3a* enhancer (cyan) positioned upstream of the zebrafish E1b minimal promoter (grey) and driving eGFP (green) expression harbouring either Ds- or Tol2-integration arms (black). 30 pg of either construct (“Ac/Ds” and “Tol2”) was microinjected into *Gt(FoxD3:mCherry)^ct110aR^* embryos (to visualise neural tube) together with Ac or Tol2 mRNA, respectively. Live confocal imaging highlighted similar levels of neural crest cell labelling between 30 pg Ac/Ds and 150 pg Tol2, but 30 pg Tol2 injections yielded much weaker signal in comparison. **B** Labelling efficiencies of the Ac/Ds and Tol2 *pax3a* enhancer were quantified using immunohistochemistry and Imaris. Ac/Ds resulted in up to 9-fold increase (Student *t*-test;*p* =0.001) of GFP^+^/Hoechst^+^ signal compared to Tol2. **C** The same approach in (**B**) was used to compare labelling efficiency of a *sox10* enhancer with weak activity. Although both versions resulted in comparable GFP^+^/Hoechst^+^ signal, Ac/Ds demonstrated less variability compared to Tol2. Scale bar: 40 µm.

These results demonstrated a clear advantage of Ac/Ds- over Tol2-mediated enhancer testing, and in general transient expression systems using DNA constructs in zebrafish, as Ac/Ds transposition yielded a higher number of construct integrations resulting in lower cell mosaicity rate. This was particularly notable with a strong element such as the *pax3a* enhancer, where a high number of cells showing consistent enhancer activity pattern were successfully labelled. Furthermore, the utility of Ac/Ds in being able to inject both DNA and mRNA at lower amounts is significant. Toxicity issues associated with the higher levels required for Tol2-based somatic integration could be avoided, and lower amounts of DNA required for high integration efficiency minimises ectopic activity from episomal expression of non-integrated plasmid. This places the zebrafish embryo on par with the chick embryo as an excellent model for testing enhancer activity transiently in injected F_0_ embryos (Streit et al., 2013). Taken together, we demonstrated that Ac/Ds-mediated integration is not only more efficient for transgenes with strong activity but at the same time more efficacious for transgenes with weaker activity.

### *In vivo* characterisation of novel neural crest enhancers using Ac/Ds

Having established the utility of Ac/Ds for testing enhancers in F_0_ embryos, we further characterised the *pax3a* and *sox10* enhancers described earlier following germline transmission (Fig.2, *pax3a* E5 and *sox10* E2). We found that germline transmission rates for Ac/Ds-injected embryos were comparable to Tol2, consistent with previous reports (Emelyanov et al., 2006, Emelyanov and Parinov, 2008). We also characterised three additional enhancers - one predicted for *pax3a* and two for *sox10* (Fig.2; *pax3a* E4, *sox10* E5 and E7) in both F_0_ and following germline transmission (*pax3a* E4), or in F_0_ only (*sox10* E5 and E7) (Supp.Fig.3). *Pax3a* and *sox10* are well-characterised transcription factors with known roles in neural crest development (Alkobtawi et al., 2018). *Pax3a* E4 and E5 are located 23.5 and 6kb upstream of the transcription start site (TSS) of *pax3a* at open chromatin regions detected by ATAC-seq (Buenrostro et al., 2013) performed on FAC-sorted neural crest cells (Trinh et al., 2017, Gavriouchkina et al., 2017) (Fig.2A, maroon track). *Sox10* E2, E5 and E7 are located 33, 13kb upstream of the TSS and 3.7kb downstream of the 3’UTR of *sox10*, respectively (Fig.2B, maroon track).

**Figure 2.**
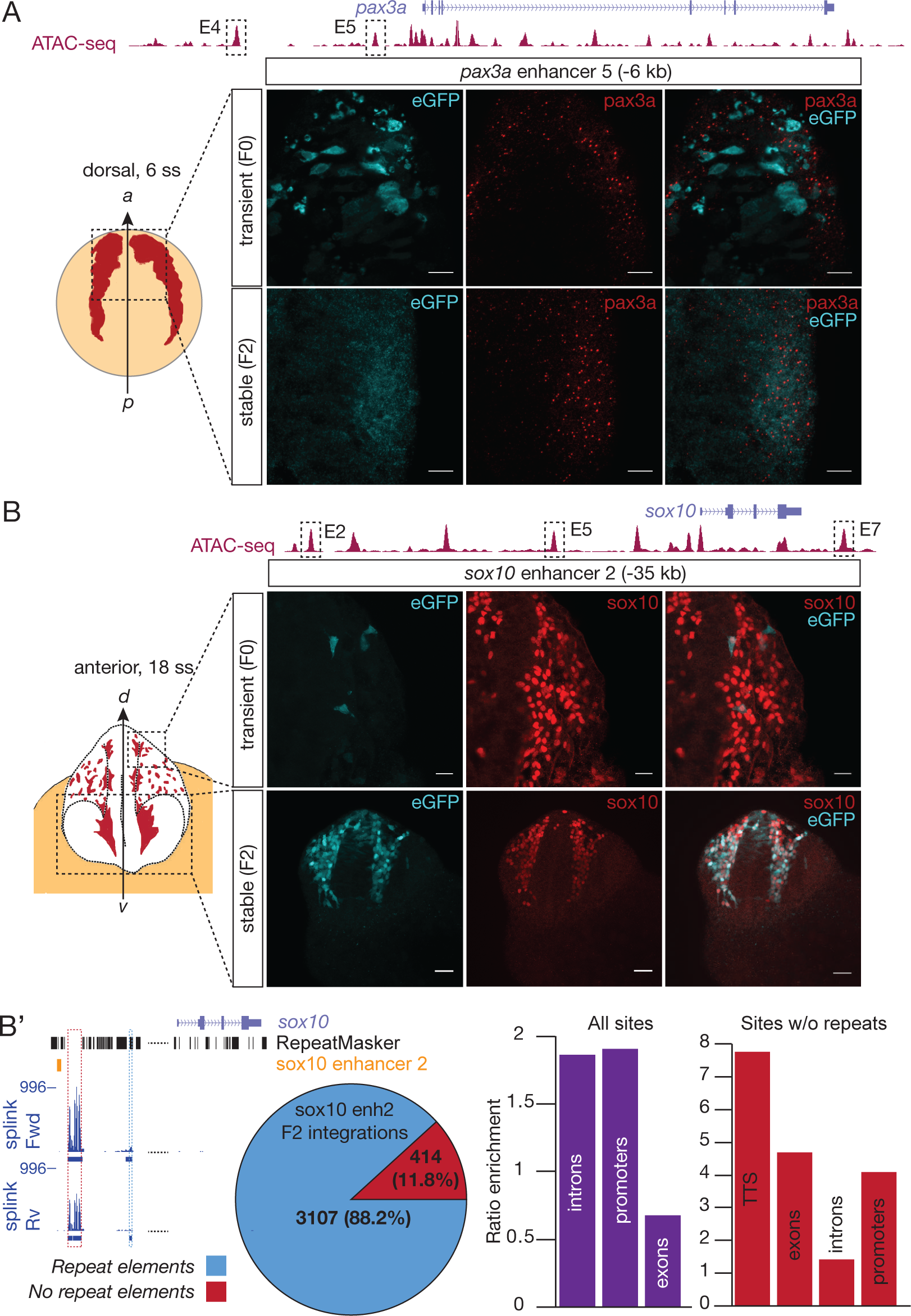
*In vivo* characterisation of *pax3a* and *sox10* enhancers using Ac/Ds integration. Full legend on next page. **A** Two putative *pax3a* enhancers (E4 and E5; E5 also shown in Fig 1) are visualised on UCSC Genome Browser (maroon, ATAC-seq track; indigo, *pax3a* gene locus). E5-driven eGFP transcripts in F_0_ and F_2_ embryos demonstrated a similar expression pattern that overlapped endogenous *pax3a* mRNA. Note the post-fixing eGFP protein signal detected in F_0_ embryo. **B** Three putative *sox10* enhancers (E2, E5 and E7; E2 also shown in Fig 1) are visualised on UCSC Genome Browser (maroon, ATAC-seq track; indigo, *sox10* gene locus). EGFP and endogenous Sox10 proteins were detected using immunohistochemistry. Weak E2-driven eGFP activity labelled very few Sox10-positive cells in F_0_ embryos. F_2_ embryos demonstrated a remarkably higher number of Sox10-positive cells with eGFP activity. **B’** Integration sites of the *sox10* E2:eGFP transgene in F_2_ embryos identified using splinkerette-PCR-NGS. Reads corresponding to flanking genomic regions on Ds-3’ end (navy blue; splink Fwd) and Ds-5’ end (navy blue; splink Rv) were mapped to the zebrafish genome (GRCz10) and peaks corresponding to integration sites were bioinformatically called (navy blue boxes). The *sox10* locus is shown as an example, with the position of *sox10* E2 highlighted in orange. 88.2% of integration sites partially/fully overlapped annotated repeat elements (blue; highlighted with dotted box), while the remaining 11.8% (red; highlighted with dotted box) did not. Annotated introns, promoters and exons were enriched (*p<*0.04) within all integration sites (purple bar chart). Transcription termination sites (TTS) were also enriched (*p<*0.01) within integration sites that did not overlap annotated repeat elements (red bar chart). Scale bar: 20 µm; 40 µm (**B**-stable (F2)).

To test the co-localisation of *pax3a* expression with *pax3a* E4 or *pax3a* E5 activity, we utilised the Hybridisation Chain Reaction (HCR) method (Choi et al., 2018) to detect *eGFP* and endogenous *pax3a* mRNAs. At 6 somite stage (ss), *pax3a* is strongly expressed in premigratory neural crest at the neural plate border region (Fig.2A, cartoon reproduced from Zfin *in situ* data). Co-localisation of *eGFP* and *pax3a* mRNA transcripts in this region were detected in F_0_-injected embryos, and this result was recapitulated in F_2_ embryos, *Tg(pax3a enh5-E1b:eGFP)^ox163^* (Fig.2A). *Pax3a* E4 gave a similar result but with overall weaker eGFP expression in both F_0_ and F_2_, *Tg(pax3a enh4-E1b:eGFP)^ox162^* (Supp.Fig.3).

Next, we tested the co-localisation of Sox10 expression with *sox10* E2, E5 and E7 activity using immunohistochemistry to detect eGFP and endogenous Sox10 proteins. At 18 ss, *sox10* is strongly expressed in migratory neural crest within the cranial region (Fig.2B, cartoon reproduced from (Gavriouchkina et al., 2017)). All *sox10* enhancers demonstrated weak but Sox10-specific activity in F_0_-injected embryos (Fig.2B, Supp.Fig.3). Remarkably, *sox10* E2-driven eGFP expression dramatically improved following germline transmission with a much higher number of GFP^+^/Sox10^+^ cells being detected in *Tg(sox10 enh2-E1b:eGFP)^ox120^* F_2_ embryos (Fig.2B). Interestingly, this “germline-boosting” effect was not observed in five other enhancer lines that we have propagated to the germline.

To recover integration sites of the transgene in a genome-wide fashion, we performed splinkerette-PCR-NGS on a pool of GFP^+^ F_2_ embryos (Supp.Fig.4, Materials & Methods). High-confidence sites were defined as genomic regions where the signal of mapped reads from splinkerette PCR amplicons representing genomic regions flanking Ds-5’ and 3’ arms were enriched over background (qval<0.01) (Fig.2B’, splink Rv and splink Fwd tracks). We identified 3,521 high-confidence sites, significantly higher than a previous study reporting 1,685 integration sites across 424 Ac/Ds transgenic lines (Vrljicak et al., 2016). We reasoned that our result may represent most of the initial integration events still present in the F_1_ germline, as indicated by the 5,473 putative inserts identified in F_0_-injected embryos by Vrljicak *et al.*. Echoing the previous study, we also found a large proportion of integration sites (88.2%) to overlap annotated repeat elements (Fig.2B’, pie chart) and enriched at gene regions with broad distribution across promoters, introns, exons and transcription termination sites (TTS) (Fig.2B’, red and purple bar chart) (Vrljicak et al., 2016).

The results demonstrated by the weak *sox10* E2 enhancer fortuitously highlighted the potential for our approach to uncover candidates that would otherwise have been overlooked at the F_0_ screening stage if Tol2-integration was used. The higher likelihood of a false positive result with Tol2 could influence a negative decision for the propagation of the transgene as a reporter line, thus eliminating potential lines that could be biologically relevant. In short, we showed that the Ac/Ds enhancer construct is a robust and useful tool to characterise putative enhancers with both strong and weak activity.

### Ac/Ds successfully expresses sgRNAs for transient tissue-specific CRISPR*i*

The ability to knockdown non-coding elements *in vivo* is essential for the dissection of their function. Currently, the most commonly used method to deliver sgRNAs for CRISPR/Cas9 in zebrafish is via microinjection of *in vitro*-transcribed sgRNAs into single cell embryos. While cost-effective and straightforward, this approach risks decreasing the efficiency for CRISPR-mediated events in the embryo, as the uncapped and non-polyadenylated nature of sgRNAs renders them sensitive to degradation *in vivo* (Mir et al., 2018). This is particularly pertinent if the desired goal is to perform CRISPR experiments in a tissue-specific fashion, as unprotected sgRNAs injected in the absence of Cas9 protein or Cas9 mRNA are likely to be diminished by the time a tissue-specific Cas9 protein is expressed many hours later in development.

We took advantage of Ac/Ds to generate a constitutive sgRNA expression system that employs a “transient transgenesis” approach for use in tissue-specific CRISPR/dCas9 (nuclease-deficient Cas9) modulation of targeted non-coding loci and enhancers in F_0_ embryos. To express sgRNAs, we cloned a cassette containing a zebrafish-specific U6a promoter driving expression of sgRNAs (Yin et al., 2015) into a custom-made Ac/Ds mini-vector (Fig.3A, Materials & Methods). The 20bp spacer region within the cassette was flanked by *Bsm* BI restriction sites to facilitate GoldenGate (Clarke et al., 2012)-like cloning of different sgRNAs (Supp.Fig.5A). We compared continuous expression of a scrambled sgRNA sequence from the integrated Ac/Ds-U6a:sgRNA vector (pVC-Ds-DrU6a:sgRNA-Ds) and an *in vitro*-transcribed sgRNA transcript by injecting 50 pg of vector (with 24 pg *Ac* mRNA) or 80 pg of transcript into wild type embryos, followed by RT-PCR at 5 hours post injection (hpi), 24 hpi and 5 days post injection (dpi). We found that sgRNA expression up to 5 dpi could only be detected using Ac/Ds (Fig.3A). To achieve tissue-specific (neural crest) expression of dCas9 fused to repressive effector protein (SID4x) for CRISPR/dCas9-interference (CRISPR*i*) at target regions (Fig.3B), we generated BAC transgenic line *TgBAC(sox10:dCas9-SID4x-2a-Citrine)^ox117^* (“ox117”) by taking advantage of the previously validated *sox10* BAC clone (Trinh et al., 2017). This approach enabled neural crest-specific expression of dCas9-SID4x in an endogenous *sox10*-like fashion, under the control of regulatory elements embedded within the *sox10* regulatory locus contained in the BAC clone (Fig.3C).

**Figure 3.**
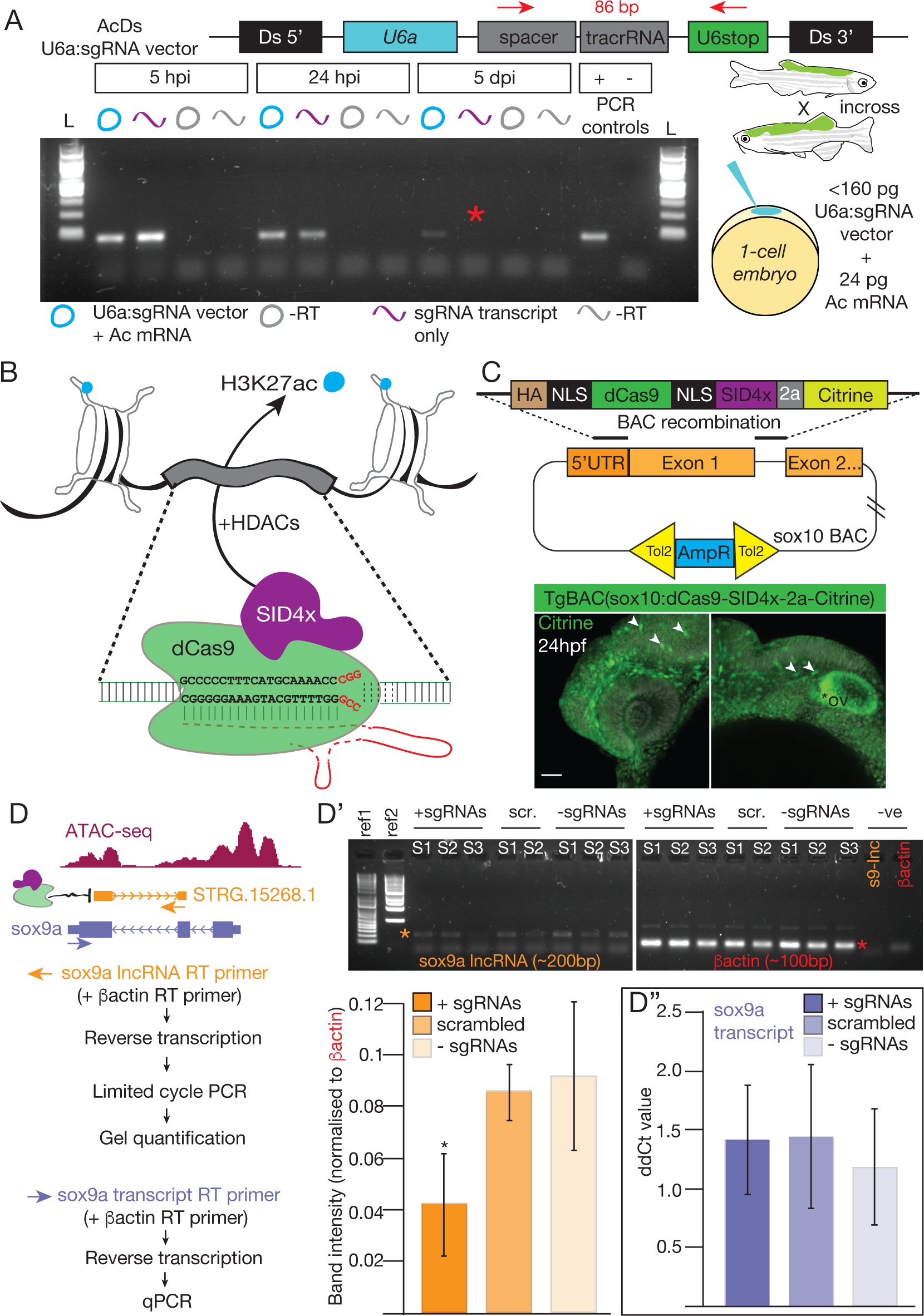
Ac/Ds ubiquitous expression of sgRNAs for Sox10-specific CRISPR*i* in the neural crest of transgenic F_0_ embryos. Full legend on next page. **A** A zebrafish U6a promoter is used to drive expression of an sgRNA cassette consisting of spacer region (20bp target sequence), tracrRNA and a U6 termination sequence (U6stop). The entire transgene is flanked by Ds integration arms (black). AcDs-U6a:sgRNA was microinjected into incrossed ox117 embryos and its sgRNA transcript detected by RT-PCR (red arrows indicate primers; product size 86bp) at 5 hours post-injection (hpi), 24 hpi and 5 days post-injection (5 dpi). In parallel, embryos microinjected with *in vitro*-transcribed sgRNA (with same sequence as AcDs version) were also accessed. At 5 dpi, sgRNA expression was only detected in embryos microinjected with AcDs-U6a:sgRNA. **B** Nuclease-deficient Cas9 (green; dCas9) fused to the SID4x repressor domain (violet) is targeted to a region-of-interest using sgRNAs (red). The SID4x domain enables transcriptional repression via chromatin compaction, possibly by recruiting histone deacetylases (HDACs). **C** A Sox10-specific CRISPR*i* transgenic line, *TgBAC(sox10:dCas9-SID4x-2a-Citrine)^ox117^*, was generated using BAC recombination and Tol2-mediated transgenesis. A ribosome-skipping TaV-2a peptide (2a) mediates separation of dCas9-SID4x from its Citrine reporter protein. The transgene successfully labels neural crest (white arrows) and the otic vesicle (ov)*in vivo*. **D** Five AcDs-U6a:sgRNAs targeting the TSS of STRG.15268.1, a transcript overlapping *sox9a* in the opposing strand (“*sox9a* lncRNA”), were microinjected into ox117 incrossed embryos. Strand-specific reverse transcription (RT) was performed using primers that bind specifically to the predicted 3’ end of *sox9a* lncRNA (orange) or the 3’UTR of *sox9a* mRNA (indigo). The *sox9a* mRNA RT primer is positioned upstream of *sox9a* lncRNA’s TSS target region. In both cases, *βactin* was also primed as an endogenous control.*Sox9a* lncRNA and *sox9a* mRNA were measured using semi-quantitative PCR and qPCR, respectively. **D’** Semi-quantitative PCR revealed a modest knockdown (Student *t*-test; *p*=0.03) of *sox9a* lncRNA in the presence of sgRNAs compared to controls. **D”** *Sox9a* mRNA transcript levels were unaffected by sox9a lncRNA CRISPR*i*, indicating a strand-specific mode-of-action. Scale bar: 40 µm.

We reasoned that the small size of the Ac/Ds-U6a:sgRNA mini-vector (<4.5kb), coupled with the small load required for activity, would permit pooling of multiple sgRNAs. We tested a load of up to 160 pg DNA per embryo, where the survival rate was ∼ 50% in our hands. Multiplexing sgRNAs is crucial for CRISPR*i*, given previous studies demonstrating the requirement for multiple sgRNAs to illicit successful knockdown (Qi et al., 2013, Williams et al., 2018). Previous evidence has also highlighted CRISPR*i*’s potential for strand-specific mode-of-action when dCas9 is used to target transcription elongation, but not initiation, using sgRNAs that bind the non-template strand (Qi et al., 2013). To explore this possibility in the zebrafish, we microinjected incrossed ox117 embryos with an Ac/Ds-sgRNA pool containing five sgRNAs that target the TSS of STRG.15268.1, an antisense transcript overlapping *sox9a* (“*sox9a* lncRNA”), while avoiding targeting of *sox9a* elongation (Supp.Fig.5B). The *sox9a* lncRNA was identified by *de novo* assembly (Pertea et al., 2015) of previously published neural crest RNA-seq datasets (Trinh et al., 2017) and is preceded by a strong ATAC-seq peak at its 5’ terminus, indicative of promoter activity (Buenrostro et al., 2013) (Fig.3D, maroon track).

A major obstacle to the study of lncRNAs is their significantly lower levels of expression compared to their protein-coding counterparts (Derrien et al., 2012). To circumvent this issue and quantify *sox9a* lncRNA levels following CRISPR*i*, we performed semi-quantitative transcript-specific RT-PCR with *actin* as endogenous control (Fig.3D). To quantify *sox9a* while taking into account potential strand-unspecific effects, *sox9a* was reverse-transcribed using a gene-specific primer targeted at its 3’UTR and localised upstream of the five sgRNAs targeting the TSS of *sox9a* lncRNA. Transcript levels were measured using qPCR as per standard practice (Fig.3D). We found modest knockdown (*p*=0.03) of *sox9a* lncRNA in the presence of sgRNAs compared to scrambled sgRNAs or uninjected controls (Fig.3D’), while *sox9a* transcript levels were unaffected in the same samples (Fig.3D”). These findings were consistent with our observation that injected embryos were morphologically normal and support recent evidence that lncRNAs were largely dispensable for development (Goudarzi et al., 2018).

In short, we demonstrated that re-purposing of the Ac/Ds approach enabled constitutive expression of sgRNAs in embryos *in vivo* following genomic integration of the construct into somatic cells. By pooling multiple sgRNAs, we also provided proof-of-principle evidence that CRISPR*i* could be achieved in a strand-specific fashion on a locus with overlapping antisense transcription. As an aside, we have also generated modified versions of the Ac/Ds-sgRNA vector containing RNA scaffolds for KRAB transcriptional repression (Zalatan et al., 2015) and for the synergistic transcription activation mediator (SAM) system (Konermann et al., 2015). The latter can be coupled with the *TgBAC(sox10:Cas9m4-VP64-2a-Citrine)^ox118^* line we have also generated, to enable for CRISPR/dCas9-activation (CRISPR*a*) of targeted loci (Mali et al., 2013) (Supp.Material).

### CRISPR*i* of *sox10* enhancers affects Sox10 expression

Finally, we investigated the functional relationship between *sox10* E2, E5 and E7 enhancers, characterised by our Ac/Ds enhancer assay, and endogenous sox10 expression using our optimised CRISPR*i* approach (Fig.4A), as enhancer effect on gene expression is thought to be additive (Hay et al., 2016, Will et al., 2017). To test the contributions of multiple enhancers to *sox10* gene activity, *TgBAC(sox10:dCas9-SID4x-2a-Citrine)^ox117^* incrossed embryos were injected with a pool of 15 Ac/Ds sgRNAs targeting all three *sox10* enhancers. Embryos were fixed and immunohistochemistry performed to detect Citrine (cells expressing dCas9-SID4x) and nuclei containing endogenous Sox10 protein (Materials & Methods). We identified three different scenarios in terms of co-expression - (1) dCas9-SID4x^+^ only* cells, (2) sox10^+^/dCas-SID4x^+^** double-positive cells and (3) Sox10^+^ only cells (this scenario was observed due to faint levels of Citrine at the stage of analysis) (Fig.4B). We reasoned that such a combined outcome and scenario (3) with a decrease in Citrine occurred also due to the fact our approach did not only target sox10 E2, E5 and E7 enhancers within the endogenous *sox10* locus, but also within the regulatory region included in the ox117 BAC allele driving dCas9-SID4x. As a consequence, Citrine and dCas9-SID4x expression was progressively decreased and with it lessened the effect of epigenome modulation. To quantify this phenomenon, we decided to focus on initial stages dCas9-SID4x activity to avoid the decrease in its activity (and thus release of inhibition) due to the described regulatory conundrum. Three different embryos (+/- Ac/Ds sgRNAs) were subjected to the same immunohistochemistry protocol and imaged; dCas9-SID4x^+^ cells were counted on individual slices [i.e. scenario (1)+(2)] and scenario (2), see Materials & Methods, Supp.Material). The knockdown effect was calculated as a ratio of Sox10^+^/dCas-SID4x^+^** cells in dCas9-SID4x^+^ (* + **) counted cells (Fig.4B’). Similar to the *sox9a* lncRNA knockdown experiment, we showed that deactivating these three enhancers modestly diminished Sox10 expression (*p*=0.02) (Fig.4B’).

**Figure 4.**
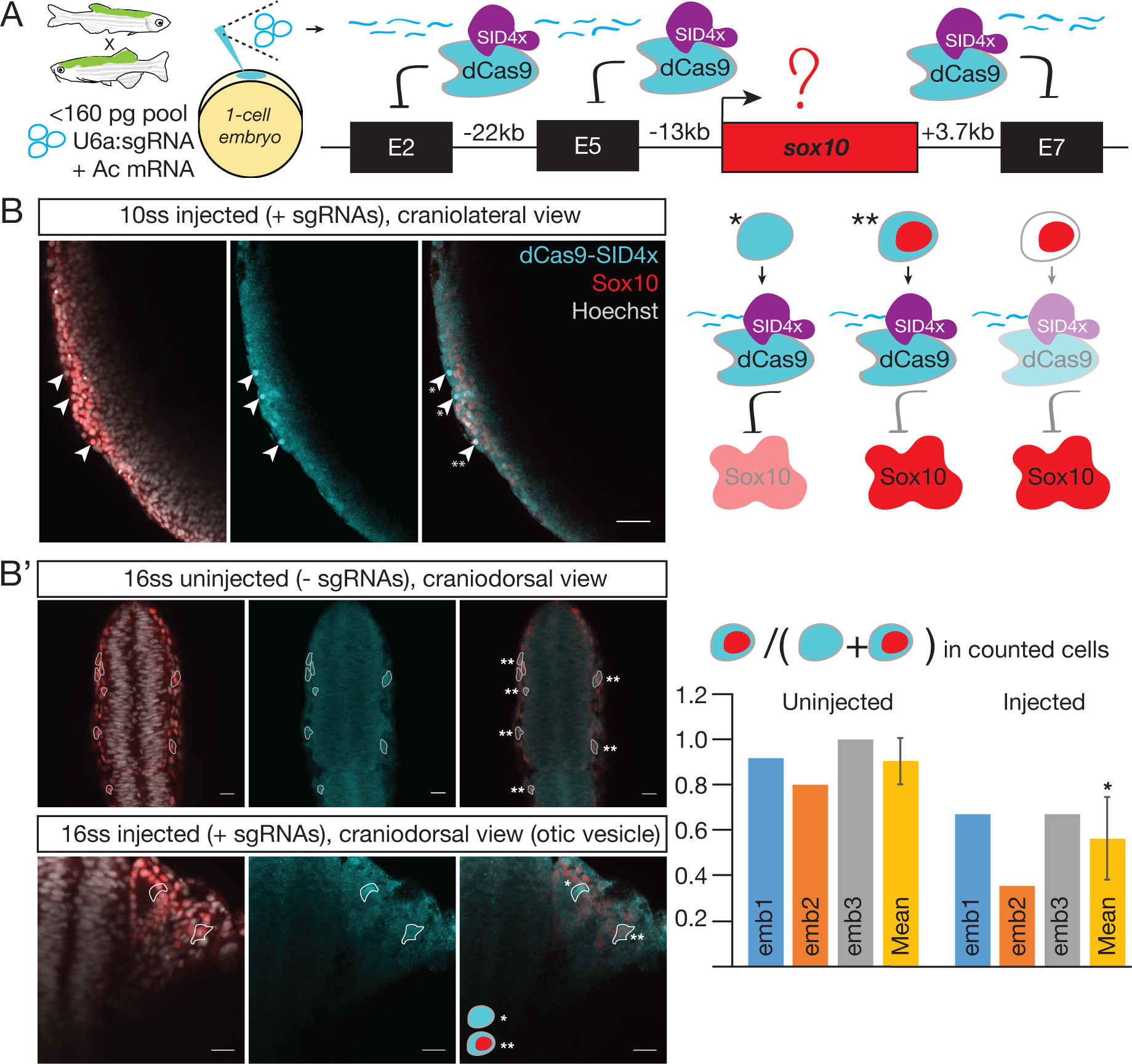
CRISPR*i* of *sox10* enhancers affects Sox10 expression. **A** CRISPR*i* was performed on three *sox10* enhancers described earlier (E2, E5 and E7) by simultaneously microinjecting five AcDs-U6a:sgRNAs per enhancer (15 in total) into ox117 incrossed embryos. Immunohistochemistry followed by confocal imaging to detect Citrine and Sox10 proteins was used as a qualitative and quantitative readout of interference. **B** Two outcomes (*; Sox10 protein undetected and **; Sox10 protein detected) were observed as a result of *sox10* E2,E5,E7 CRISPR*i* in cells where Citrine (ergo dCas9-SID4x) is present, indicating incomplete knockdown and/or secondary effects of the sgRNAs targeting *sox10* E2,5,7 that are also present on the dCas9-SID4x BAC allele. **B’** To quantify this observation, cells with high Citrine expression (dCas9-SID4x^+^) were individually counted in two separate slices per embryo across three different embryos per condition. The ratio of Sox10^+^/dCas-SID4x^+^ events in dCas9-SID4x^+^ counted cells revealed a modest decrease (Student *t*-test;*p* =0.02) in Sox10 expression following *sox10* E2,E5,E7 CRISPR*i*. Scale bar: 40 µm (**B**); 20 µm (**B’**).

In conclusion, these results demonstrated the utility of our Ac/Ds approach as an exploratory tool to cell-specifically probe enhancers in order to elucidate their function *in vivo*, without the need for or prior to laborious generation of stable transgenic lines for further characterisation.

## Materials & Methods

### 1. Zebrafish husbandry

All zebrafish experiments were conducted according to regulated procedures authorised by the UK Home Office within the framework of the Animals (Scientific Procedures) Act 1986. Wild type and transgenic embryos were derived from AB or AB/TL mix strains.

### 2. Plasmids and oligo sequences

Full lists of plasmids (including Addgene submissions) and oligo sequences are available as Supplementary Material.

### 3. Enhancer vector cloning

Putative enhancer elements were amplified from genomic DNA by PCR and cloned into pVC-Ds-E1b:eGFP-Ds (Addgene ID 102417) linearised with *Nhe*I. Cloning was performed using In-Fusion^TM^ HD Cloning Plus (Takara) or Gibson Assembly cloning (Gibson et al., 2009). Cloning reactions were transformed into chemo-competent cells and plasmids were prepared using QIAprep Spin Miniprep kit (27104, Qiagen) and stored in Elution Buffer. Inserts where verified by Sanger sequencing using T7 primer.

### 4. Cloning of Ac/Ds-U6a:sgRNA vector(s)

We previously generated a mini-vector with Ds arms flanking a multiple cloning site (pVC-Ds-MCS-DS; Addgene ID 102416). The insert consisting of zebrafish U6a promoter, spacer region, tracrRNA, and U6 termination sequence (Yin et al., 2015) was amplified by PCR using a custom-ordered gBlock Gene Fragment (Integrated DNA Technologies) as template. The PCR product was gel-purified and cloned into the mini-vector linearised with *Sna*BI and *Nhe*I, using In-Fusion^TM^ HD Cloning Plus (Takara). To clone desired sgRNA targets, complementary oligo pairs were ordered as follows: (1) TTCG-5’[20bp target without PAM]3’ and (2) AAAC-5’[20bp target without PAM in reverse complement]3’ (Supp.Material). 50 µM of each oligo were combined in a 50 µL reaction and annealed in a thermocycler (94°C 5 mins, decrease 20 to 22°C at 1°C/min, 4°C hold). Cloning reaction was prepared by combining 70 ng pVC-Ds-DrU6a:sgRNA-Ds vector, 5 ng annealed sgRNA, 10U of *Bsm*BI and 20U of T4 DNA ligase (M0202, NEB) in 1X T4 DNA Ligase Buffer with final volume 20 µL. ‘GoldenGate’ cycling conditions were used as follows: 10X(37°C 5 mins, 16°C 10 min), 50°C 5 mins, 80°C 5 mins. 2 µL of the reaction was transformed into chemo-competent cells and plated onto Ampicillin plates. Two colonies per sgRNA were screened by Sanger sequencing using U6a promoter primer TCACTCACCACCTCCCAAAA.

### 5. *Ac* transposase mRNA synthesis

To prepare *Ac* transposase mRNA, pAC-SP6 (Addgene ID 102418) (Emelyanov et al., 2006) was linearised with *Bam*HI and purified under RNase-free conditions.*In vitro* transcription was performed using mMESSAGE mMACHINE^TM^ SP6 Transcription Kit (AM1340, ThermoFisher). mRNA was purified under RNase-free conditions using phenol-chloroform followed by ethanol precipitation and the pellet resuspended in RNase-free water. mRNA quality was assessed by gel electrophoresis (sharp intact band without degradation) and quantified using Qubit^TM^ RNA HS Assay kit (Q32852, ThermoFisher). For long term storage in −80°C the purified mRNA was prepared as 1 µL aliquots and limited to one freeze-thaw cycle. Prior to use, an aliquot is freshly diluted with nuclease-free water and the excess discarded after use.

### 6. Ac/Ds microinjections

All microinjections were performed by injecting 2.3 nL into the blastula of one-cell stage embryos within 5 to 20 mins post fertilisation. To prepare microinjection aliquots, Miniprepped plasmids in Elution Buffer were diluted at least 20-fold with nuclease-free water. The following conditions have been optimised using the Nanoject II system (Drummond Scientific) and may have to be adjusted if using a different microinjector.

#### 6.1 Enhancer screening

For Ac/Ds vectors, each embryo was injected with 30 pg of DNA and 24 pg *Ac* mRNA. To compare with Tol2 vectors, each embryo was injected with 30 pg of DNA and 80 pg *Tol2* mRNA, or 150 pg of DNA and 50 pg *Tol2* mRNA (lethality ∼ 50%).

#### 6.2 Ac/Ds-U6a:sgRNA injections for CRISPR*i*

Single sgRNA injections were performed with 50 pg of DNA and 24 pg *Ac* mRNA per embryo. For pooling more than one sgRNA, up to 160 pg of DNA and 24 pg *Ac* mRNA were injected per embryo (maximum 15 different sgRNAs).

### 7. Ac vs Tol2 quantification

Following eGFP (with Hoechst) immunohistochemistry (see Section 14), dorsal views of embryos were imaged as 50 µm Z-stacks on LSM780 Inverted confocal microscope (Zeiss) using the same laser and acquisition settings throughout. Raw images were rendered on Imaris (Bitplane) as “Spots” and “Surfaces” for Hoechst and GFP channels, respectively. To compute total GFP intensity (*x*), GFP Mean Intensity values of “Spots” with *d>* 0 (*d*=distance from Hoechst “Spot” to GFP “Surface”), was summed. GFP intensity per 1 µm^2^ “Surface” Area, *y*, was calculated by dividing *x* with total “Surface” Area detected. GFP intensity for 50 µm^2^, *y*\*50, was plotted as a bar chart.

### 8. Ac/Ds-U6a:sgRNA vector vs *in vitro-transcribed* sgRNA RT-PCR

Total RNA was extracted from pools of 11 microinjected embryos per condition (vector, or IVT) using RNAqueous-Micro Total RNA Isolation Kit (AM1931, ThermoFisher). Reverse transcription (RT) was performed using 0.5 µM each of R sgRNA scaffold tail (Supp.Material) in a 10 µL reaction (1 µg starting RNA) using SuperScript^TM^ III Reverse Transcriptase (18080093, ThermoFisher). Reverse transcription was performed at 55°C for 60 mins. Primary PCR was performed using the following primers: R sgRNA and F scrambled1 spacer (Supp.Material). In a 20 µL reaction, 0.1 µM per primer was combined with 1 µL of template (reverse transcription reaction) in 1X standard *Taq* polymerase PCR reaction. Cycling was performed as follows: 95°C 5 mins, 35X(95°C 30s, 55°C 30s), 68°C 30s, 12°C hold. Next, secondary PCR was performed using the following primers: R nested sgRNA and F nested scrambled1 spacer (Supp.Material). In a 20 µL reaction, 0.1 µM per primer was combined with 1 µL of template (primary PCR reaction) in 1X standard *Taq* polymerase PCR reaction. Cycling was performed as follows: 95°C 5 mins, 23X(95°C 30s, 55°C 30s), 68°C 30s, 22 12°C hold. Results were analysed on a single 2% agarose gel using a 100bp ladder.

### 9. Generation of CRISPR transgenic lines

*TgBAC(sox10:dCas9-SID4x-2a-Citrine)^ox117^* and *TgBAC(sox10:Cas9m4-VP64-2a-Citrine)^ox118^* (Mali et al., 2013) were generated using BAC recombination followed by Tol2 transgenesis as previously described (Suster et al., 2011, Trinh et al., 2017). Recombination cassettes were amplified from pGEM-T-Easy-HA-NLS-dCas9-NLS-SID4x-TaV-2a-Citrine-FRT-Kan-FRT and pGEM-T-Easy-HA-NLS-Cas9m4-NLS-VP64-TaV-2a-Citrine-FRT-Kan-FRT (Supp.Material) to introduce 50bp homology arms for recombination into the *sox10* locus in BAC clone DKEY-201F15, by replacing *sox10*’s first exon with the recombination cassette.

### 10. Sox10 quantification following *sox10* E2-7 CRISPR*i*

Following Sox10 and Citrine immunohistochemistry (see section 14), Z-stack images of the anterior craniolateral region were obtained using 2.5 µm slices. Two slices (at least 15 µm apart) per embryo were used for analysis. First, cells (or cell clusters) with strongest anti-GFP (Citrine) signal (ergo dCas9-SID4x) were highlighted. Next, the highlighted GFP signal was superimposed onto the Hoechst channel to refine dCas9-SID4x^+^ cell count. Finally, dCas9-SID4x^+^/Hoechst^+^ cells with Sox10 signal were counted. This process was repeated for all slices used in the analysis.

### 11. *sox9a* lncRNA RT-PCR

Total RNA was extracted from pools of 10 CRISPR*i*-microinjected embryos per condition (+sgRNAs, +scrambled sgRNAs, -sgRNAs) using RNAqueous-Micro Total RNA Isolation Kit (AM1931, ThermoFisher).*Sox9a* lncRNA-specific reverse transcription (RT) was performed using 0.5 µM each of R STRG.15268.1, R2 STRG.15268.1 and bactin E4 R3 (Supp.Material) in a 10 µL reaction (maximising amount of starting RNA) using SuperScript^TM^ III Reverse Transcriptase (18080093, ThermoFisher). Reverse transcription was performed at 55°C for 60 mins.*Sox9a* lncRNA and βactin PCR were performed using the following primers: R2 STRG.15268.1 and R sox9aE3; beta-actin E3 and beta-actin E4 (Supp.Material). In a 20 µL reaction, 0.2 µM per primer was combined with a 1 µL of template (1:10 dilution of reverse transcription reaction) in 1X Phusion^R^ High-Fidelity PCR Master Mix with HF Buffer (M0531, NEB). Cycling was performed as follows: 98°C 5 mins, 23X(98°C 30s, 57°C 30s), 72°C 30s, 12°C hold. Results were analysed on a single 1.5% agarose gel using the 100bp band of 1kb or 1kb Plus ladder (NEB) as reference. Relative band intensities to selected reference were quantified using BioRad’s Image Lab^TM^ Software.

### 12. *sox9a* qPCR

Both the total RNA extracted for *sox9a* lncRNA RT-PCR as well as the RT protocol were re-utilised for *sox9a*-specific reverse transcription. Instead, RT primer combination used were R sox9a 3UTR and bactin E4 R3. QPCR was performed using the Ct method with 300 nM of *sox9a* primers (F sox9a E2 and R sox9a E3) and 150 nM *actin* primers (beta-actin E3 and beta-actin E4) in Fast SYBR^TM^ Green Master Mix (4385612, ThermoFisher). 0.5 µL of template (1:10 dilution of reverse transcription reaction) was used in a 10 µL reaction.

### 13. Splinkerette-PCR-NGS

#### 13.1 Preparation of splinkerette library

To assess a wide spectrum of possible integration sites of the Ac/Ds *sox10* enhancer 2 transgene, Tg(sox10 enh2-E1b:eGFP)^ox120^ F1 individuals were outcrossed and 25 GFP^+^ F2 embryos were collected for genomic DNA extraction. As a control, genomic DNA was also extracted from wild type embryos and processed in parallel. 500 ng of genomic DNA was digested with 20U of *Alu* I (R0137, NEB) in a 30 µL reaction at 37°C in a thermocycler with heated lid for 4 hours. The digest reaction was purified using phenol-chloroform extraction followed by ethanol precipitation and the pellet resuspended with 34 µL of nuclease-free water. Splinkerette adaptors (SPLINK-top VC and SPLINK-bottom VC; Supp.Material) were annealed at 25 µM final concentration each in 2.5 µL volume with the following cycling conditions: 95°C 2 mins, decrease 95 - 22°C at 0.1°C/second and ending with 22°C 5 mins before placing on ice. The ligation reaction was prepared by combining 17 µL of purified *Alu* I-digested genomic DNA with 0.5 µL annealed adaptors and 10U of T4 DNA ligase (M0202, NEB) in 1X T4 DNA ligase buffer. Ligation was incubated 24 overnight at 16°C in a thermocycler with heated lid, subsequently purified using DNA Clean & Concentrator (D4003, Zymo Research) and eluted with 25 µL DNA Elution Buffer. A primary PCR reaction (12.5 µL volume) was setup by combining 10 ng of ligated DNA with 0.25 µL each of 10 µM SPLINK P1 and DS-3’ forward or DS-5’ reverse P1 primers (Supp.Material) in 1X Platinum SuperFi master mix (12358010, ThermoFisher). Cycling was performed as follows: 98°C 2 mins, 25X(98°C 10s, 63°C 30s, 72°C 3 mins), 72°C 5 mins, 12°C hold. After cycling, 1 µL of the primary PCR reaction was combined with 1 µL each of 10 µM SPLINK P2 and DS-3’ forward or DS-5’ reverse P2 primers (Supp.Material) in 1X Platinum SuperFi master mix (50 µL volume) for secondary PCR. Cycling was performed like primary PCR, with the exception that annealing temperature was lowered to 60°C. 5 µL of the secondary PCR reaction was run on a 2% agarose gel to assess amplification of genomic regions flanking integrations. Prior to library prep for Next Generation Sequencing (NGS), the secondary PCR reactions were purified using Select-a-Size DNA Clean & Concentrator (D4080, Zymo Research) according to manufacturer’s instructions to enrich for amplicons between 100-1000bp. Purified amplicons were quantified by Qubit^TM^ dsDNA HS Assay kit (Q32854, ThermoFisher) and 50 ng used as starting input for library prep. Library prep was performed using Nextera DNA Library Prep Kit (FC-121-1030, Illumina) according to manufacturer’s instructions and their fragment profiles (∼200bp expected) assessed on the TapeStation (Agilent Technologies). Libraries were sequenced on NextSeq500 Illumina platform (v2 150 cycles, 80-bp paired end) to obtain ∼ 4 million reads per forward/reverse sample.

#### 13.2 Bioinformatics analysis

Obtained reads were quality trimmed using sickle (Joshi and Fass, 2011) and -l 30 -q 30 parameters. Trimmed reads were mapped to the *D.rerio* genome (GRCz10) using STAR (Dobin et al., 2013) with default parameters. Book-ended mapped regions on the same strand were merged using bedtools merge (Quinlan and Hall, 2010)) to ‘recapitulate’ amplicon fragments prior to tagmentation during library prep. To identify regions with significant signal over background in mapped reads (10 to 50-fold enrichment, ergo integration sites), peaks were called using MACS2 callpeaks (Zhang et al., 2008) with -g 1.41e9 -m 10 50 --nomodel --shiftsize X -q 0.01 parameters, where X=library 25 fragment size/2. To filter out ‘high-confidence’ called peaks, bedtools intersect was used to retrieve common peaks found in both forward and reverse samples overlapping by at least 1bp. If desired, peaks that overlapped annotated repeat elements by at least 1bp were removed using bedtools subtract. Genome ontology analysis of peaks were performed using HOMER annotatePeaks.pl (Heinz et al., 2010) with `-genomeOntology intersect_positions_genomeOntology-gsize` 1.41e9 parameters.

### 14. Whole-mount immunohistochemistry

Embryos were fixed in 4% paraformaldehyde (PFA) for 45 minutes at room temperature (RT). After washing, embryos were blocked using 10% goat serum in PBT (PBS, 0.5% Triton, 2% DMSO) for 1 hour at RT. Primary antibodies used were chicken anti-GFP (to detect eGFP or Citrine) (ab13970, Abcam) and rabbit anti-zfSox10 (GTX128374, GeneTex) in a 1:200 dilution, added overnight at 4°C. Secondary antibodies used were donkey anti-rabbit 568nm (A10042, ThermoFisher) and goat anti-chicken 647nm (A21449,ThermoFisher) in 1:400 dilution each, together with Hoechst reagent (H3569, ThermoFisher) in a 1:1000 dilution to label nuclei - all added for 2 hours at RT. After washing to remove excess, embryos were imaged using a LSM780 confocal microscope (Zeiss).

### 15. Hybridisation Chain Reaction

Embryos were fixed in 4% paraformaldehyde (PFA) overnight at 4°C, washed in phosphate-buffered saline (PBS), dehydrated in methanol (MeOH) and stored at −20°C. Hybridization chain reaction (HCR) v3.0 was performed according to published protocol (Choi et al., 2018). Briefly, embryos were rehydrated with a series of graded MeOH/PBS-Triton (PBST) washes and incubated overnight at 37°C in 30% probe hybridization buffer containing 2 pmol of each probe mixture (pax3a and GFP). Excess probes were washed off with 30% probe wash buffer at 37°C and 5XSSCT at RT. Embryos were then incubated overnight at RT in amplification buffer containing 15 pmol of each fluorophore-labelled hairpin (B3-546 and B2-488). Excess hairpins were removed by washing with 5XSSCT at RT. Following HCR, embryos were imaged using a LSM780 confocal microscope (Zeiss). 26

## Author contributions

Conceptualisation, V.C.-M., T.S.-S.; Methodology, V.C.-M., U.S., F.C.S.; Investigation, V.C.-M., F.C.S., D.S.C.; Writing - Original Draft, V.C.-M.; Writing - Review & Editing, all authors; Supervision, T.S.-S.; Funding Acquisition, T.S.-S., P.R.R.

## Acknowledgements and funding

This work was supported by MRC (G0902418), Lister Institute Research Prize, Oxford BHF CRE award (RE/08/004) to T.S.-S. and (RE/13/1/30181) to F.C.S., RDM Pump Priming Grant (HSD00040) to T.S.-S. and V.C.-M., Clarendon Fund Fellowship to V.C.-M, BHF Programme grant and Chair award (RG/13/9/303269 and CH/11/1/28798) to P.R.R.. We would like to thank Wenbiao Chen for advice on choice of zebrafish U6a promoter, Sarah de Val for E1b minimal promoter cassette, Joey Riepsaame for dCas9-SID4x construct, and current collaborators who have requested the enhancer construct for their work.

